# Nutraceutical and functional potential of the *Apis mellifera* L. royal pupae proteome

**DOI:** 10.64898/2026.03.11.709969

**Authors:** Delcer Guadalupe Ruíz-Pérez, Roberto De la Rosa-Santamaría, Itzel López-Rosas, Said Cadena-Villegas, Jenner Rodas-Trejo, Francisco Izquierdo-Reyes, Luis Manuel Vargas-Villamil

## Abstract

The objective of this research was to characterize the proteome of the *Apis mellifera* royal pupae to evaluate its potential as a nutraceutical and functional food. Six pupal instars (E1–E6) were analyzed using liquid chromatography, mass spectrometry, and bioinformatics techniques to determine their properties and biological functions.

The results showed 15 proteins across the different instars. In E1, the Isoform X2 of the Caf1 protein and the vitellogenin precursor were found, both critical in genetic regulation and nutrient transport. E2 revealed three proteins linked to energy and genetic processes. Proteins identified in E3 were associated with sugar metabolism and cellular structure. E4 presented proteins related to cellular stress and oxidative processes. In E5, three proteins were identified, associated with molecular transport and energy metabolism. Results for instar E6 were inconclusive since the complexity of peptide identification.

From a nutraceutical and functional perspective, the identified proteins show significant potential due to their antioxidant activities, metabolic control, and cellular regulation. Noteworthy proteins include aldose reductase for its role in diabetes management, glutamate dehydrogenase for its importance in amino acid metabolism, vitellogenin as a nutrient source and immune system stimulant, and heat shock protein 60 A, with therapeutic potential in cardiovascular diseases.

## Introduction

The production and availability of nutraceutical and functional foods is a global trend whose demand is supported by sources of information such as expert opinion, research findings, and information shared through social media (Vargas-Villamil et al., 2022). A nutraceutical and functional food produces metabolic or physiological effects that promote a healthy condition in the organism, in addition to fulfilling basic nutritional functions (Meléndez et al., 2020).

The consumption of insects has been practiced by human populations since ancient times as a source of proteins, fats, and vitamins, with approximately 1900 to 2100 species recorded worldwide (Van Huis, 2013; Kaur et al., 2023); in Mexico, at least 56 edible species exist (Cruz-Labana et al., 2018), in a practice that has received greater attention in recent decades due to the therapeutic uses to which insects have been directed (Siddiqui et al., 2023), where chitin from the exoskeleton plays an important role (Van Huis, 2013), among other components whose nature and effects remain unknown.

Different studies have been conducted to estimate the nutritional value and properties of bees per se, where it has been found that a bromatological profile exists that varies among their different stages of biological development. For example, drone larvae are rich in proteins and other nutrients that should be investigated in greater detail since they contain nutritional as well as allergenic elements (Matuszewska-Mach et al., 2024). In contrast, drone pupae possess high nutritional value, are a safe food, and may exert medicinal efficacy (Choi, 2021).

As the worker larva of *A. mellifera* approaches the pupal phase, the content of carbohydrates and fats decreases, whereas protein quantities increase; the contents of iron and zinc are considerable at all developmental stages (Ghosh et al., 2016), which favors their use for food purposes (Kim et al., 2019; Kaur et al., 2023; Siddiqui et al., 2023); however, the properties of royal pupae of the honey bee remain unknown.

The development of the queen bee is an epigenetic phenomenon, resulting from the normal expression of reproductive genes induced by a diet based on royal jelly, which is received by a larva derived from a fertilized egg, in contrast with the development of workers (Ghosh et al., 2016), where the effect of royalactin, among the main components of royal jelly, induces the differentiation of larvae originating from fertilized eggs into queens (Kamboj et al., 2023).

The composition of royal jelly results from the diet received by the bees that produce it, and it contains minerals such as potassium, iron, magnesium, manganese, and phosphorus (Ghosh et al., 2024), 10-hydroxydecenoic acid (10-HDA), amino acids and proteins, among other components. This allows the hypothesis that protein content is considerable and variable in the different pupal stages of the queen of *A. mellifera*, with nutraceutical and functional properties that could be used for human consumption. The objective of the present research was to determine the proteomic profile of instars 1 to 6 of royal pupae of *A. mellifera*, and their potential as a nutraceutical and functional food.

## Materials and Methods

A descriptive observational analysis (Manterola and Otzen, 2014) was conducted on queen pupal stages of *Apis mellifera* L., during the period from 10 to 15 days of their development within the queen cell (Büchler et al., 2024).

Queen bee pupae were obtained using the Doolittle method (Büchler et al., 2024) in an apiary located in Huimanguillo, Tabasco, Mexico, situated at 18°00’ north latitude and 93°30’ west longitude, at an altitude of 9 m above sea level. The phases of the rearing method used are described in general terms (Büchler et al., 2024):

The queen’s oviposition was recorded by marking combs in the brood nest. The eggs were allowed to hatch, which occurs 72 hours after oviposition.

The starter colony was prepared one day before grafting.

Larval grafting was performed into wax queen cups, ensuring that the larvae used were of similar age, but not older than 24 hours.

The pupae contained in the queen cells were removed from the colony when they reached the appropriate age for the study.

To obtain pupae of known age, the date of grafting and the capping date of the queen cells were recorded to follow their development according to what was described by Büchler et al. (2024).

Sample collection of pupae was carried out from the moment of sealing of the queen cells. In total, six samples were taken during the pupal developmental stage. Two pupae were collected for each developmental stage, which were identified as samples E1 (10 days), E2 (11 days), successively until E6 (15 days). These were preserved in 20% trichloroacetic acid (TCA) and refrigerated at 4°C until processing for proteomic analysis (Maldonado et al., 2017).

The samples were washed seven times with a 100 mM Tris-Cl solution (pH 8.5) to eliminate TCA residues. The pupae were suspended in 200 µl of 0.1% RapiGest SF solution (Waters), and cell lysis was induced by sonication on ice using 5 pulses (30% amplitude) of 30 s each, with an interval of 2 min between pulses. The lysates were clarified by centrifugation (15,000 × g for 30 min) at 4°C. RapiGest SF is an anionic surfactant that accelerates the production of peptides in solution generated by proteases such as trypsin,

Asp-N, Glu-C, Lys-C, and PNGase F; it facilitates the digestion of proteolytically resistant proteins and increases the solubility of hydrophobic proteins and peptides. In addition, it is a mild protein denaturant compatible with mass spectrometry that helps solubilize and unfold proteins (Rapi Gest SF Surfactant Care and Use Manual, 2015).

The Pierce Micro BCA protein assay kit (Thermo Fisher Scientific) was used to quantify proteins in each sample using a standard curve with concentrations of 0, 25, 50, 125, and 250 µg of bovine serum albumin (BSA), respectively. To each concentration, 150 µl of BCA reagent was added (Reagent A 25 parts: sodium carbonate, sodium bicarbonate, sodium tartrate in 0.2 N NaOH; Reagent B 24 parts: 4% BCA in water; and Reagent C 1 part: 4% cupric sulfate pentahydrate in water). Sample absorbance was determined by measuring at a wavelength of 568 nm, plotting the data by linear regression to determine the exact protein quantities in each sample. After quantification, 100 µg of protein from each sample were used for analysis by liquid chromatography/mass spectrometry (LC-MS/MS) (Aksenov et al., 2017; Seem et al., 2021).

To identify the peptides recovered by LC-MS/MS, with the reference numbers obtained in the MASCOT database (2024), a search was conducted in the NCBI (2024), UniProt (2023), and Hymenopteramine (2023) databases for sequence analysis, including data on the size, weight, and function of each protein identified in the different pupal stages.

NCBI and UniProt are free repositories containing biological information such as genomic and protein sequences; therefore, they are selected for the analysis of nucleotide and amino acid sequences to assign identity to data from unknown or target samples. UniProt is considered the leading resource for the storage of protein data; additionally, information on the cellular localization and tertiary structure of the identified proteins was retrieved from this database.

## Results

Among the six pupal stages or instars (Ei, i = 1, 2, …, 6) analyzed (Figure 1), 15 main proteins were identified, with a range of biological, structural, enzymatic, and transport functions.

**Figure 1.**
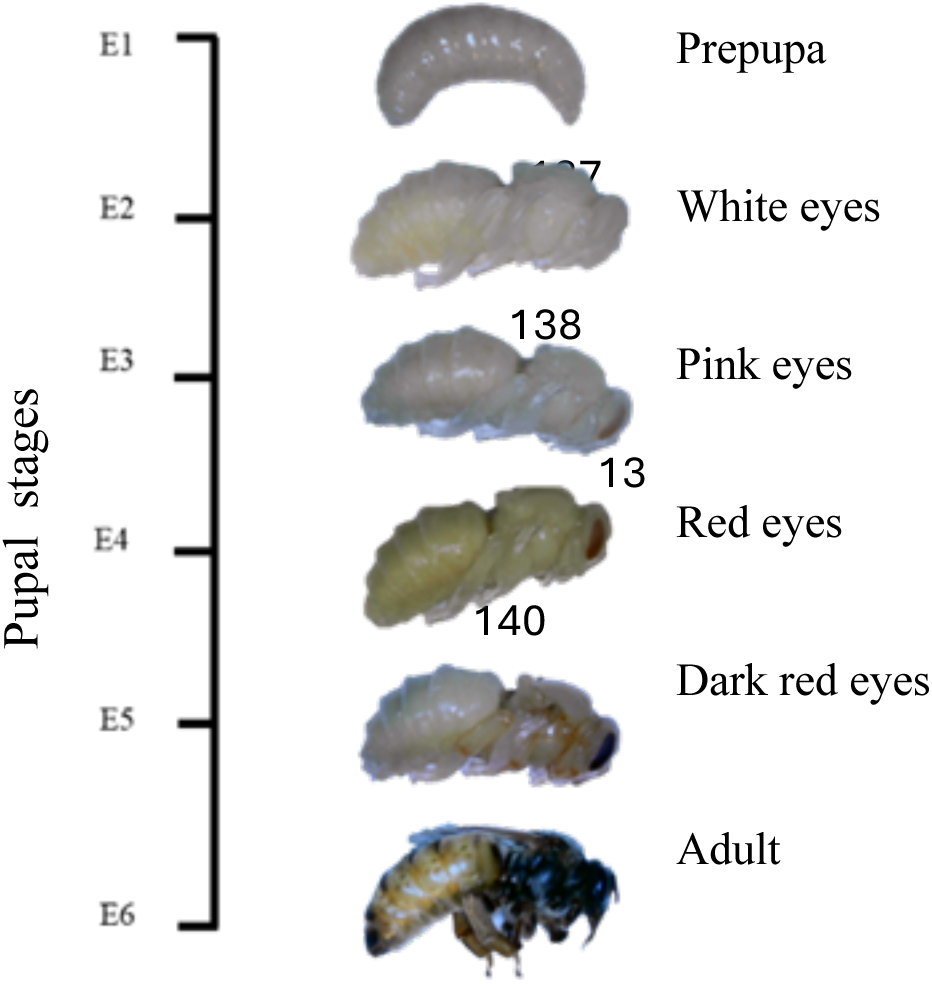
Images of the queen bee pupa of *Apis mellifera* L. at different stages (E*i*) of development. The numbers indicate the days after capping of the cells. Images without scale. Photos: Delcer Guadalupe Ruiz Pérez.

In the stage E1, two proteins were identified: Isoform X2 of the probable histone-binding protein Caf1 and a vitellogenin precursor. The first (Figure 2) has a length of 427 amino acids and a molecular weight of 48.1 kDa, and the vitellogenin precursor (Figure 3) is composed of 1770 amino acids and has a molecular weight of 201.1 kDa (Uniprot, 2023).

**Figure 2.**
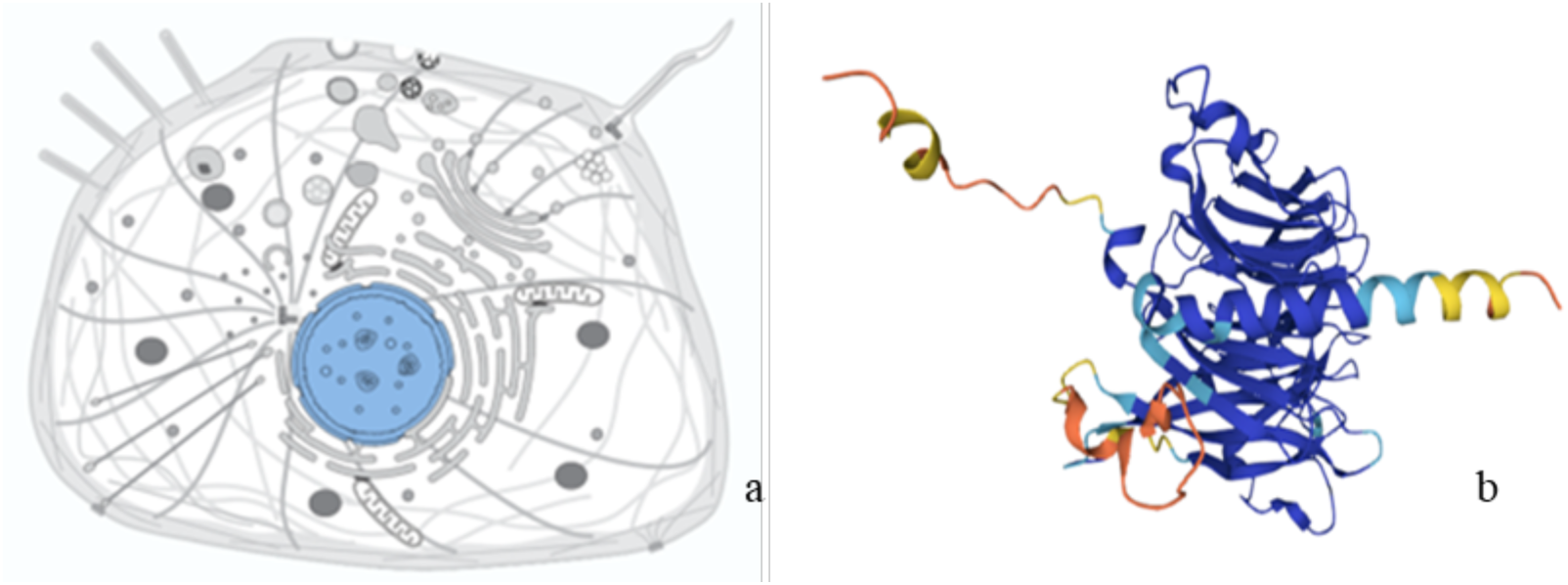
a) Localization of the probable histone-binding protein Caf1 isoform X2 in the cell nucleus, and b) Tridimensional structure. Source: UniProt, 2023.

**Figure 3.**
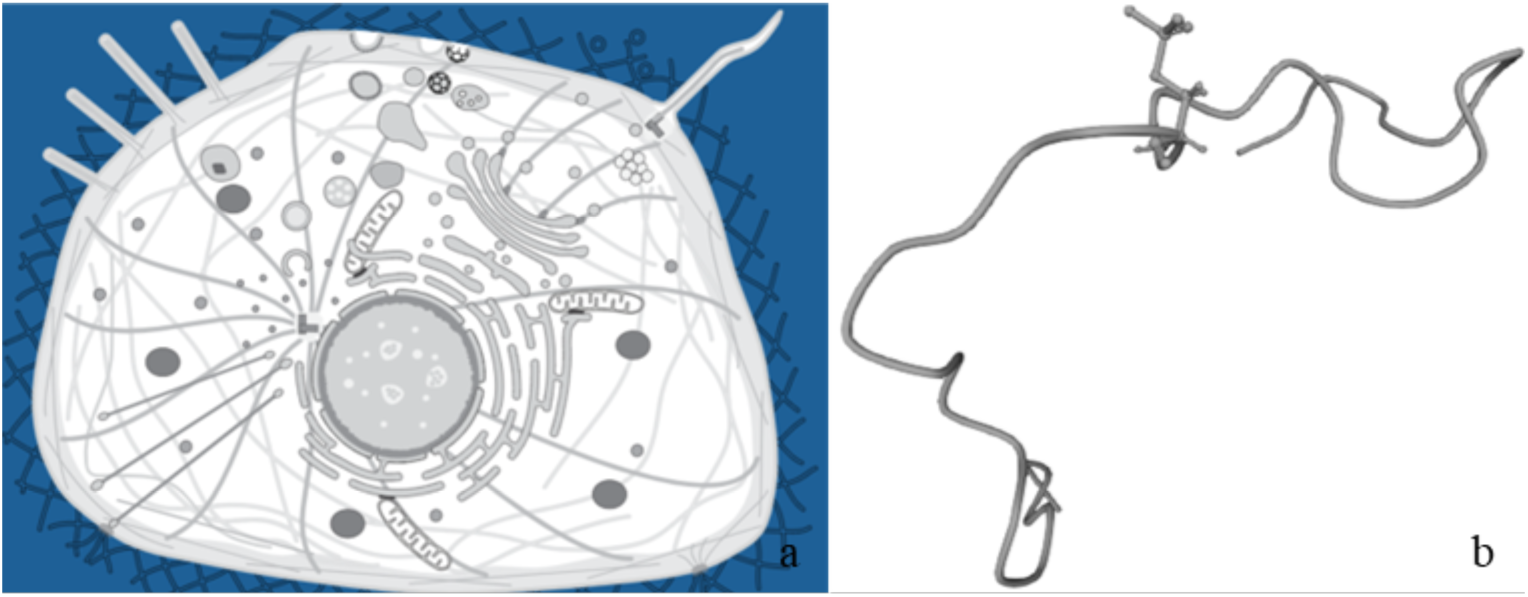
a) Localization of the vitellogenin precursor in the cell, and b) Tridimensional structure. Source: UniProt, 2023.

In the stage E2, three proteins were identified: the ubiquitin-conjugating enzyme E2 N, with a length of 151 amino acids and 17.2 kDa (Figure 4); the electron transfer flavoprotein-ubiquinone oxidoreductase, with 606 amino acids and 67 kDa (Figure 5); and pre-mRNA processing factor 19 (Figure 6) (Uniprot, 2023).

**Figure 4.**
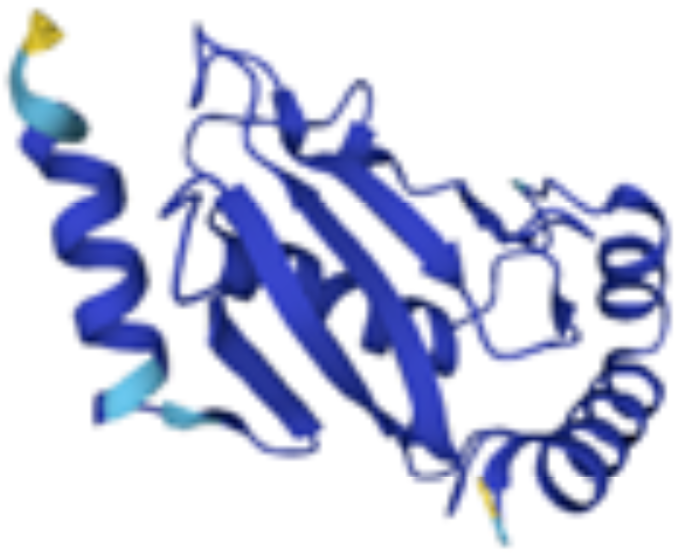
Tridimensional structure of the ubiquitin-conjugating enzyme E2 N. Source: UniProt, 2023.

**Figure 5.**
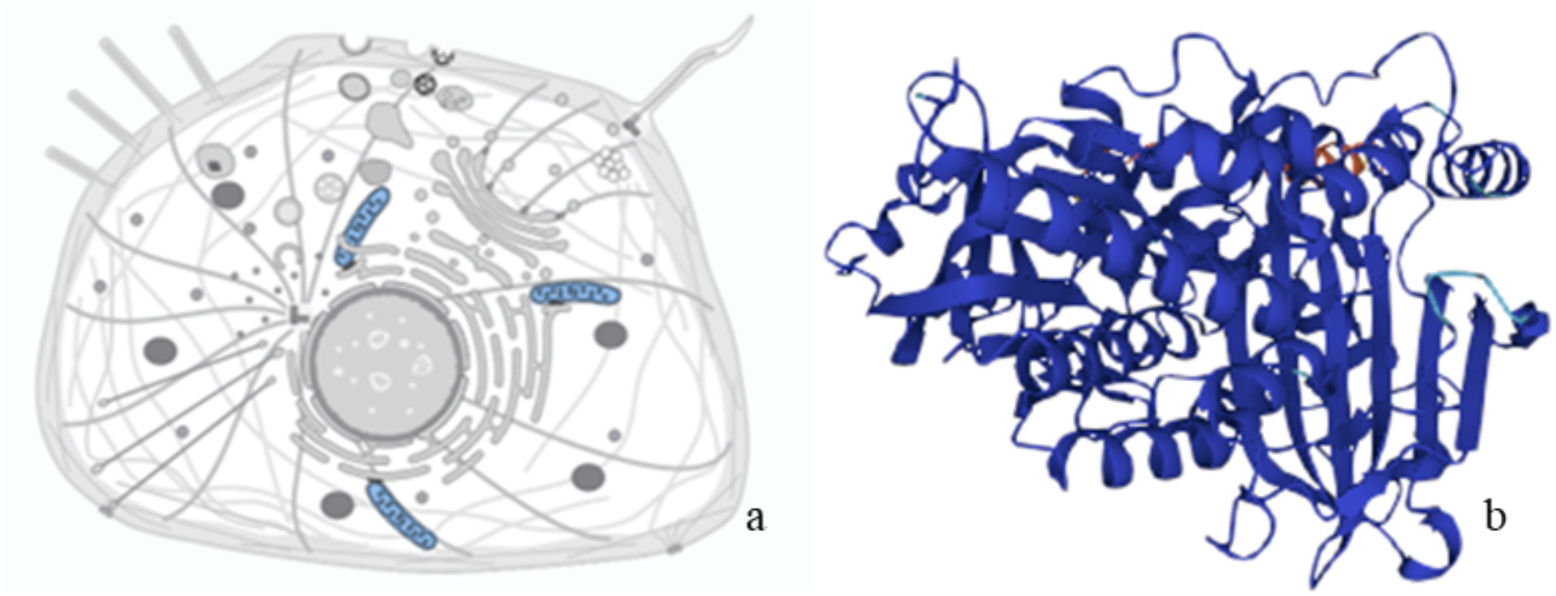
a) Localization of the mitochondrial electron transfer flavoprotein-ubiquinone oxidoreductase, and b) Tridimensional structure. Source: UniProt, 2023.

**Figure 6.**
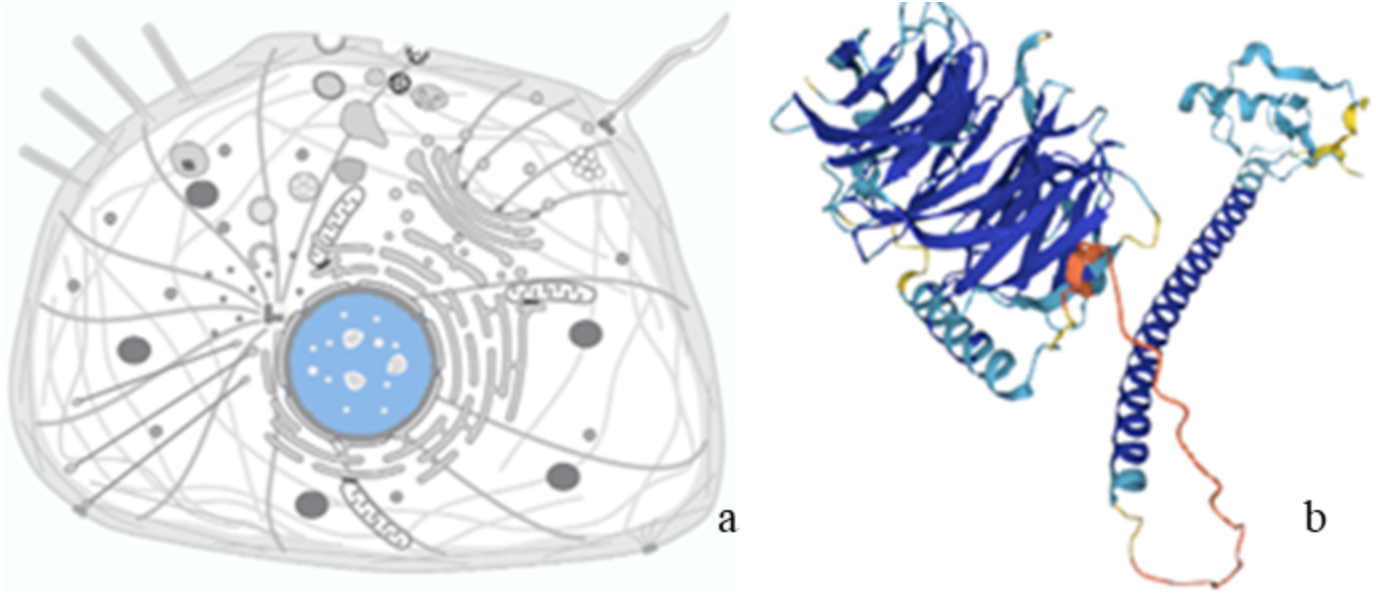
a) Localization of the pre-mRNA-processing factor 19 in the cell nucleus, and b) Tridimensional structure. Source: UniProt, 2023.

In the stage E3, three proteins were identified: an aldose reductase, a predicted protein similar to the nucleosome remodeling factor 38 kDa CG4634-PA (Figure 7), and dynactin subunit 2 (Figure 8). Aldose reductase has a length of 318 amino acids and a molecular weight of 36 kDa (Singh et al., 2021; UniProt, 2023). The predicted protein similar to the nucleosome remodeling factor −38 kDa (Figure 9) has a length of 476 amino acids and a molecular weight of 53.9 kDa (Becker and Workman, 2013; UniProt, 2023), and the dynactin subunit 2 has a length of 405 aminoacids and 46 kDa (Uniprot, 2023).

**Figure 7.**
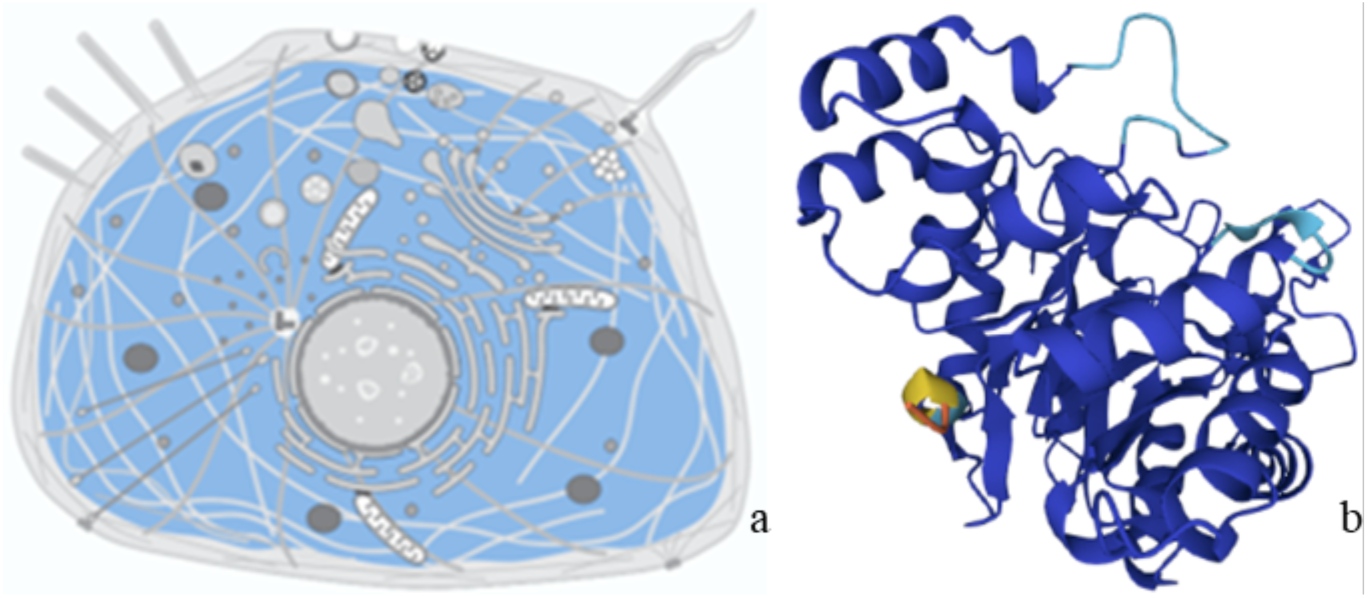
Localization of the aldose reductase in the cell cytosol, and b) Tridimensional structure. Source: UniProt, 2023.

**Figure 8.**
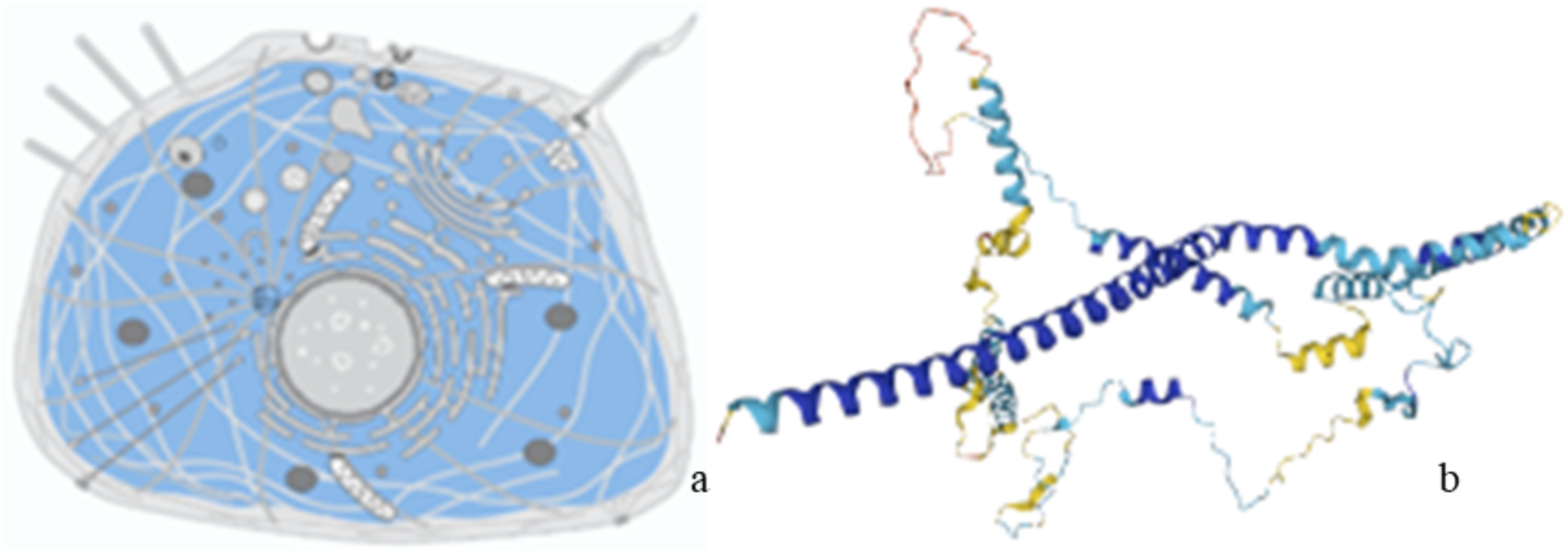
a) Localization of the dynactin subunit 2 in the centrosome-cytoplasm cell, and b) Tridimensional structure. Source: UniProt, 2023.

**Figure 9.**
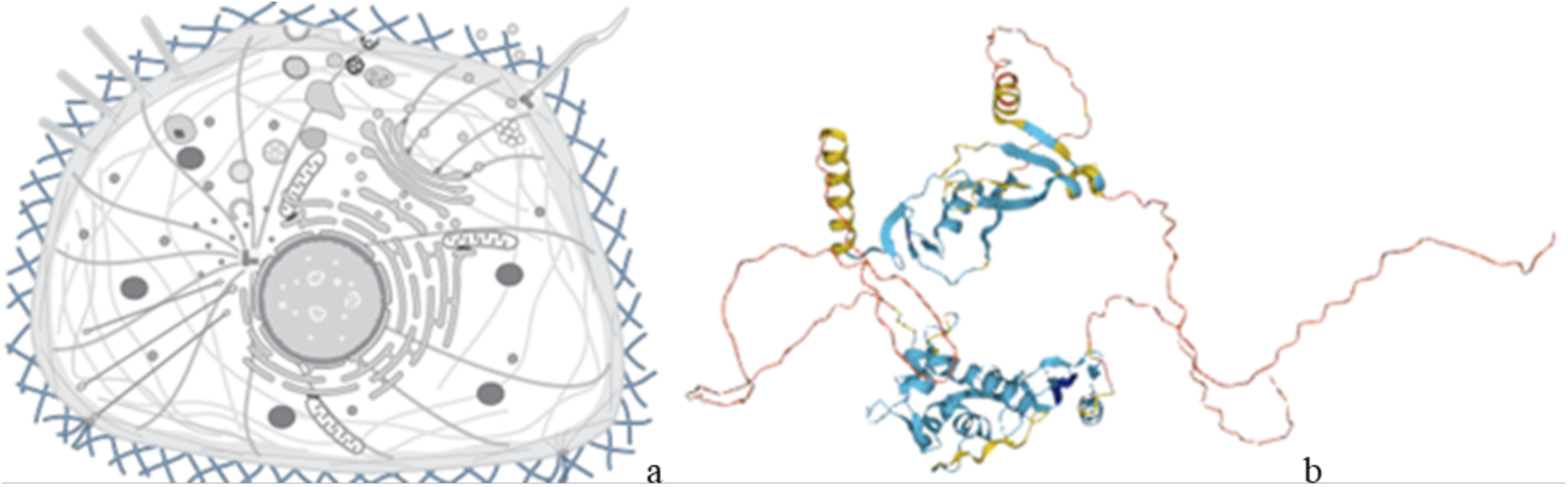
Localization of predicted similar to nucleosome remodeling factor - 38kD CG4634-PA in the extracellular matrix, and b) Tridimensional structure. Source: UniProt, 2023.

**Figure 10.**
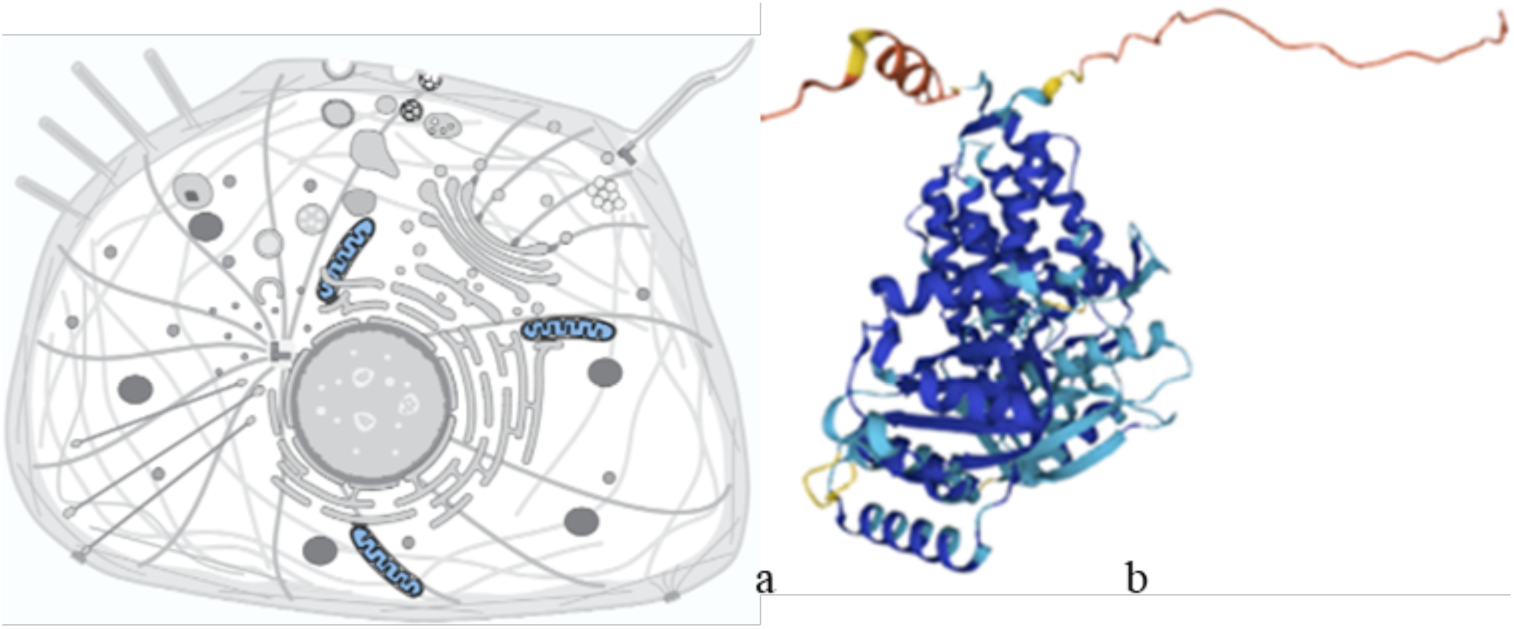
a) Localization of the heat shock protein 60A in the cell mitochondrial internal membrane, and b) Tridimensional structure. Source: UniProt, 2023.

In the stage E4, heat shock protein 60 A was found (Supplement Figure 10), with a length of 570 amino acids and a molecular weight of 60.4 kDa; thioredoxin reductase (Figure 11), with 485 amino acids in length and a molecular weight of 53.2 kDa; and isoform 2 of thioredoxin reductase 1 (Figure 12), with 494 amino acids in length and a molecular weight of 54.2 kDa (UniProt, 2023).

**Figure 11.**
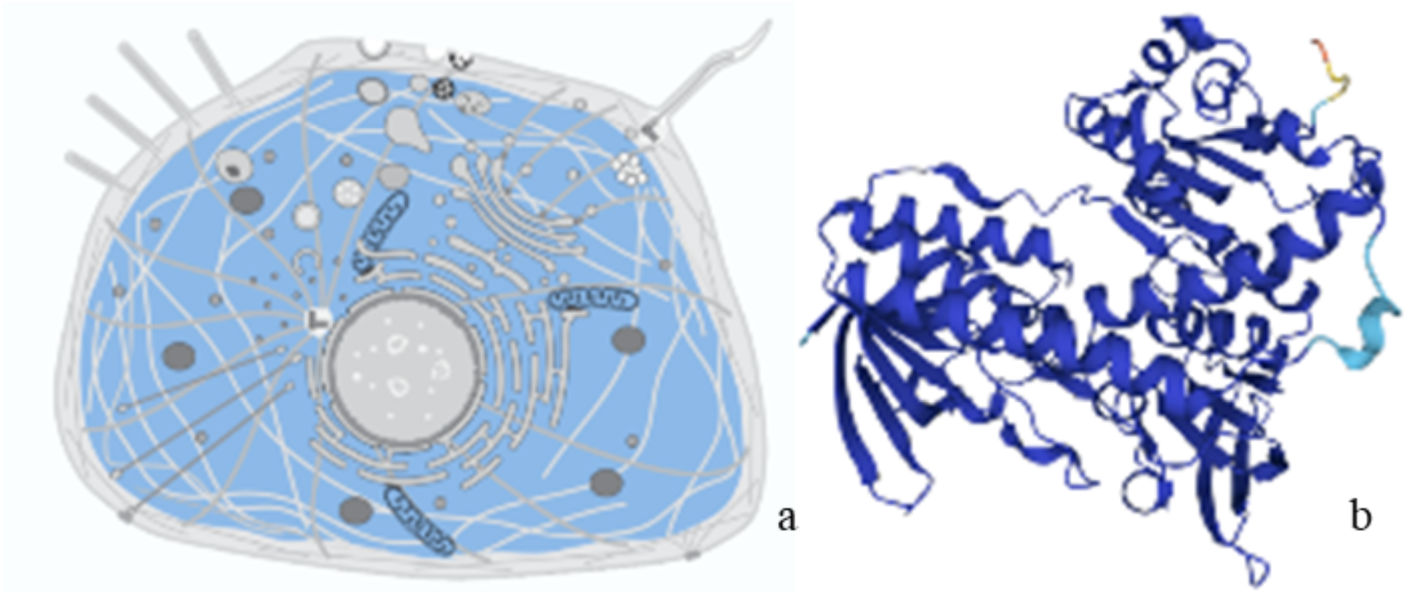
a) Localization of the thioredoxin reductase in the mitochondrial cytosol, and b) Tridimensional structure. Source: UniProt, 2023.

**Figure 12.**
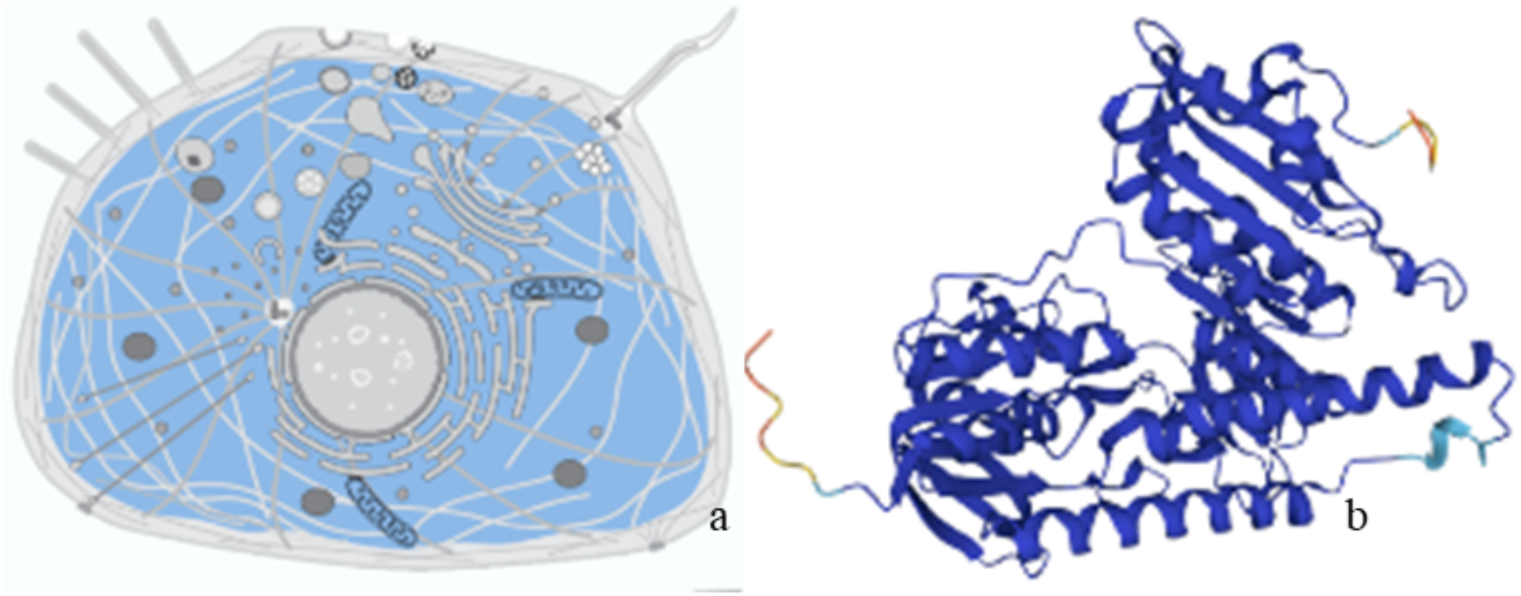
a) Localization of the thioredoxin reductase 1 isoform 2 in the cell: mitochondrial cytosol, and b) Tridimensional structure. Source: UniProt, 2023.

In the stage E5, four proteins were identified; however, only three of them have been described: the NTF2-related export protein (Figure 13), the probable medium-chain specific acyl-CoA dehydrogenase (Figure 14), and glutamate dehydrogenase mitochondrial isoform X2 (Supplement Figure 14). The export protein has 130 amino acids in length and a molecular weight of 14.7 kDa; the probable medium-chain specific acyl-CoA dehydrogenase has a length of 417 amino acids and a molecular weight of 46.5 kDa (Hymenopteramine, 2023; UniProt, 2023; Sayers et al., 2024). The mitochondrial isoform X2 of glutamate dehydrogenase has a length of 456 amino acids and a molecular weight of 45.8 kDa (Hymenopteramine, 2023; UniProt, 2023; Sayers et al., 2024).

**Figure 13.**
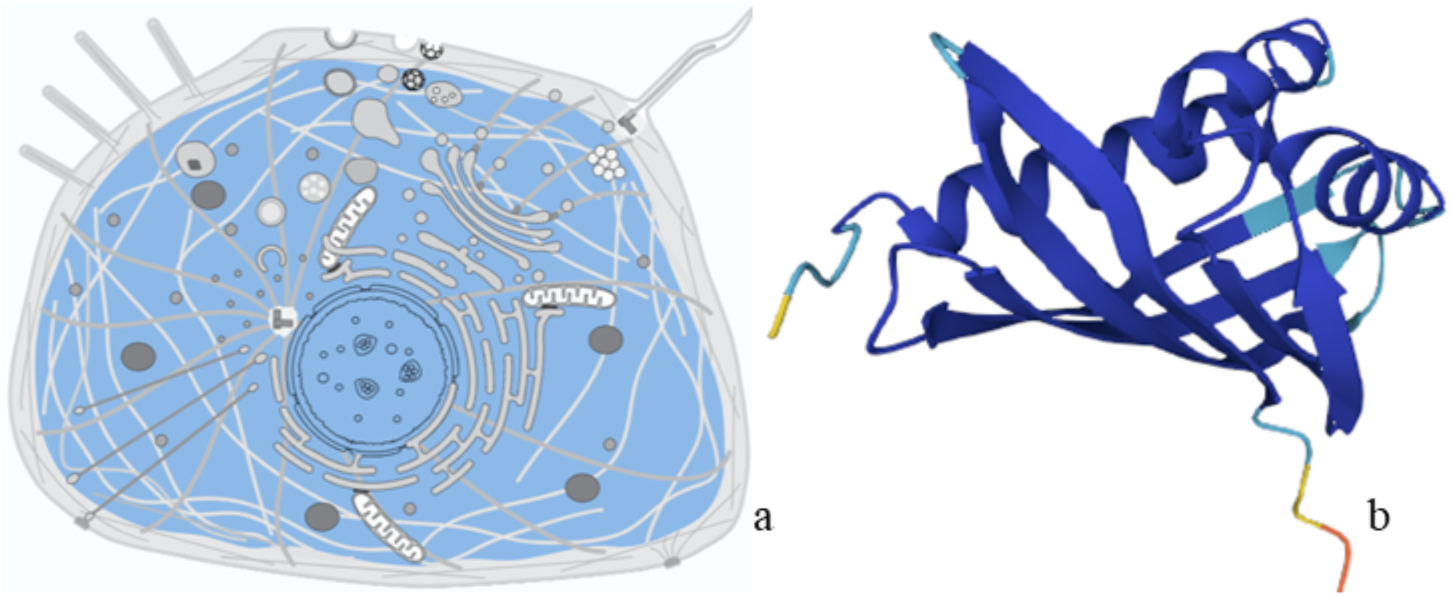
a) Localization of the NTF2-related export protein in the cell: cytoplasm and nucleus, and b) Tridimensional structure. Source: UniProt, 2023.

**Figure 14.**
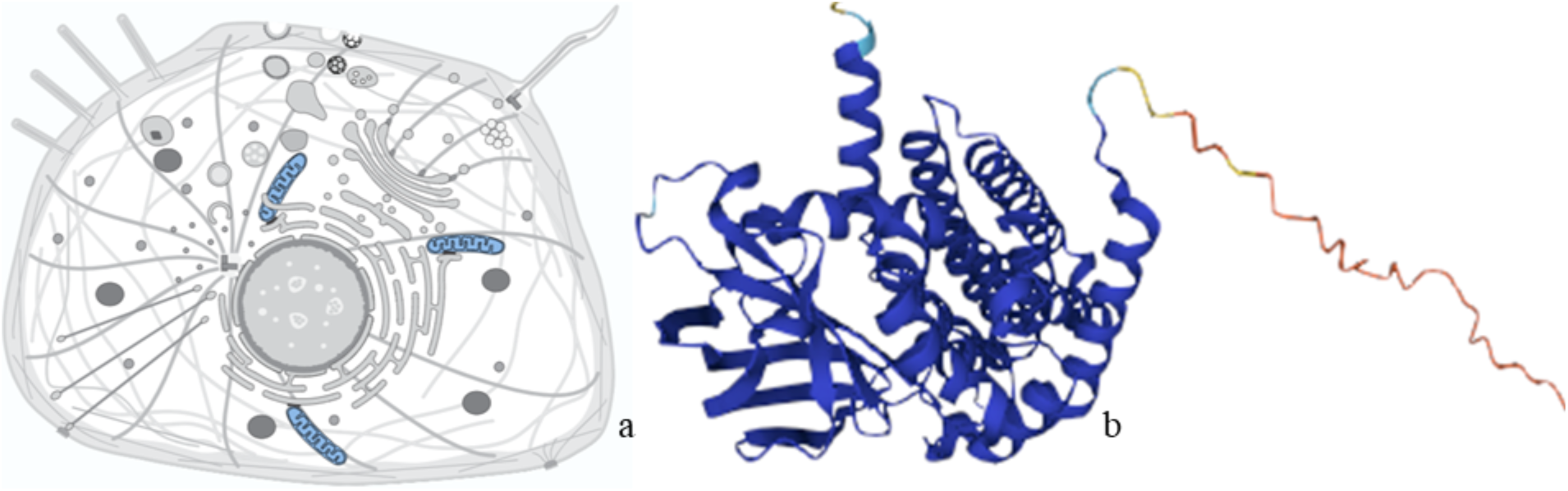
a) Localization of the mitochondrial probable medium-chain specific acyl-CoA dehydrogenase, and b) Tridimensional structure. Source: Uniprot, 2023.

The identification of peptides present in the pupal development stage E6 was not conclusive because, although computational methods allow the linking of amino acid sequences into folds of known proteins, some modules or domains belong to many different proteins, which makes it difficult to identify the particular protein to which they correspond (Alberts, 2002).

## Discussion

Some of the identified proteins are involved in key processes such as stress response, energy metabolism, cellular signaling, and gene expression (Hymenopteramine, 2023; UniProt, 2023; Sayers et al., 2024). This differs from the 58 proteins detected in honey bee workers (Zheng et al., 2011), whose synthesis is altered during stages 13 to 17, whereas in phases 19 to 20, 22 proteins were identified that are related to neuronal and hypopharyngeal gland development (Zheng et al., 2011).

Isoform X2 of the probable histone-binding protein Caf1 participates in the regulation of DNA transcription through histone binding, an action that provides accessibility to DNA and is essential for organizing genetic material in the nucleus (Rivera et al., 2016; UniProt, 2023).

The vitellogenin precursor (Figure 3) participates in lipid transport and serves as a reservoir of nutrients in the extracellular region. Its transport function contributes to the distribution of lipids where they are required, whether to build cellular membranes, store energy, or produce cellular signals (UniProt, 2023). The reservoir function allows the storage of nutrients (such as lipids, carbohydrates, and proteins) that the cell can use according to its energetic or metabolic needs, and it is essential for maintaining energy balance and supporting metabolic processes during periods of scarcity or high energy demand (Alberts et al., 2002).

Vitellogenin is also considered a phospholipoglycoprotein with the properties of a sugar, a fat, and a protein. Therefore, it represents a source of energy reserve during the early developmental stages of females of oviparous species (García et al., 2013); in *A. mellifera* its synthesis has been associated with the defense system against infections caused by *Nosema ceranae* (Antúnez et al., 2013).

The ubiquitine-conjugating enzyme E2 N (Figure 4) is responsible for the binding of ATP molecules and their transfer, providing the energy necessary for processes such as muscle contraction, molecular transport, and the synthesis of macromolecules, as well as the capacity to transfer functional groups such as phosphate, methyl, or amino groups from one molecule to another. This is fundamental in different biochemical processes, including nutrient metabolism, the regulation of cellular signaling, and protein modification (Alberts et al., 2002; Zapata et al., 2015; UniProt, 2023).

The flavoprotein–ubiquinone oxidoreductase (Figure 5) participates in the formation of a cluster composed of four iron atoms and four sulfur atoms; additionally, through electron transfer it functions as a dehydrogenase and participates in metal ion binding. These functions help in the oxidation of different molecules and provide structural stabilization (Watmough and Frerman, 2010; UniProt, 2023).

Pre-mRNA processing factor 19 in the nucleoplasm (Figure 6) participates in the removal of introns and the joining of exons in pre-mRNA; additionally, it is associated with and participates in ubiquitin ligation, DNA repair, mRNA segment binding, and K63-linked ubiquitination (UniProt, 2023). The ligase function catalyzes the binding of the ubiquitin molecule to target proteins through the marking of proteins for degradation, modification, or regulation in various cellular processes. The repair function is a process through which cells identify and correct damage in their DNA, which is crucial for maintaining genetic integrity and preventing mutations. For its part, K63-linked ubiquitination is associated with cellular signaling processes and DNA repair rather than protein degradation (Alberts et al., 2022; Yin et al., 2012).

Aldose reductase (Figure 7) plays an important role in the metabolism of sugars in the cytosol, catalyzing the conversion of aldoses, such as glucose, into their corresponding alcohols such as sorbitol, using NADPH (nicotinamide adenine dinucleotide phosphate in its reduced form) as a cofactor (Singh et al., 2021; UniProt, 2023).

Dynactin subunit 2, which is part of the dynactin complex in the cytoplasm and centrosomes (Figure 8), is a key component in the assembly and functioning of cellular transport, as well as in genetic stability and the prevention of errors in chromosome distribution that could lead to genetic anomalies. During mitosis, this function occurs through the organization of the mitotic spindle, which is composed of microtubules and regulatory proteins and is fundamental for separating duplicated chromosomes, ensuring that each daughter cell receives a complete copy of the genetic material (Wang et al., 2018; UniProt, 2023).

The predicted protein similar to nucleosome remodeling factor −38 kDa CG4634-PA (Figure 9) corresponds to a protein complex in the extracellular space that modifies chromatin structure, facilitating or restricting the access of other proteins to DNA, and plays a key role in the regulation of transcription, replication, and DNA repair. The designation CG4634-PA is a specific identifier for the predicted protein in certain genomic databases that refers to a particular organism. Its mechanism of action occurs through binding to calcium ions for cellular signaling, to collagen for tissue repair and maintenance, and to the extracellular matrix for cellular communication, adhesion, and maintenance of tissue structure (Becker and Workman, 2013; UniProt, 2023).

Heat shock protein 60 A (Figure 10) acts as a chaperone in the binding and arrangement of ATP molecules and ATP-dependent proteins. It also participates in changes related to programmed cell death and in the response to protein unfolding, both in mitochondria, as well as in the import of proteins into the spaces between mitochondrial membranes and in the refolding of other peptides. This process allows denatured or misfolded proteins to recover their native structure, facilitated by chaperones to maintain proper cellular function (UniProt, 2023).

Thioredoxin reductase (Figure 11) participates in flavin adenine dinucleotide binding, in glutathione disulfide reductase activity, and in thioredoxin disulfide reductase activity, both through the action of NADPH used as an electron donor. In this way, this protein is also related to cellular redox homeostasis, the cellular response to oxidative stress, and the metabolic process of glutathione (Mustachich and Powis, 2000; UniProt, 2023).

Isoform 2 of thioredoxin reductase 1 (Figure 12) participates in functions similar to those of thioredoxin reductase in the cytosol and mitochondria, although different isoforms may have slightly different cellular localizations or functions (UniProt, 2023). This protein fulfills three biological and three molecular functions corresponding to flavin adenine dinucleotide binding, glutathione-disulfide reductase activity (NADPH), thioredoxin-disulfide reductase activity (NADPH), cellular redox homeostasis, cellular response to oxidative stress, and the glutathione metabolic process (Arvizu et al., 2012; Vázquez, 2018).

The NTF2-related export protein facilitates the transport of molecules between the nucleus and the cytoplasm of the cell (Figure 13), and its biological functions include the transport of mRNA from the nucleus, where it is transcribed, to the cytoplasm where it will be translated into proteins, an essential step for gene expression (UniProt, 2023). Nucleocytoplasmic transport allows the movement of molecules between the nucleus and the cytoplasm of the cell, regulated by nuclear pores, and is essential for functions such as gene regulation and cellular response, as well as protein transport involving the movement of proteins between different cellular compartments such as the cytoplasm and nucleus or toward the cell membrane and organelles; this process is key for proper protein localization in the cell (Geydan et al., 2010).

The probable mitochondrial medium-chain specific acyl-CoA dehydrogenase (Figure 14) likely participates in the oxidation of medium-chain fatty acids, which contain between 6 and 12 carbon atoms, acting in the first stage of β-oxidation that converts fatty acids into energy by releasing ATP in the mitochondria. The role played by this protein is crucial in lipid metabolism and is especially important in tissues with high energy demands such as the liver and muscles (Alberts et al., 2002).

The mitochondrial isoform X2 of glutamate dehydrogenase (Figure 15) is an enzyme that uses NAD+ as a cofactor to catalyze the conversion of glutamate into α-ketoglutarate; this process is important in the Krebs cycle and in the regulation of glutamate and ammonium levels in the cell. This catabolic process favors metabolism and neurotransmission through the derivation of products that contribute to the balance of amino acids and energy in the cell (Plaitakis et al., 2017).

**Figure 15.**
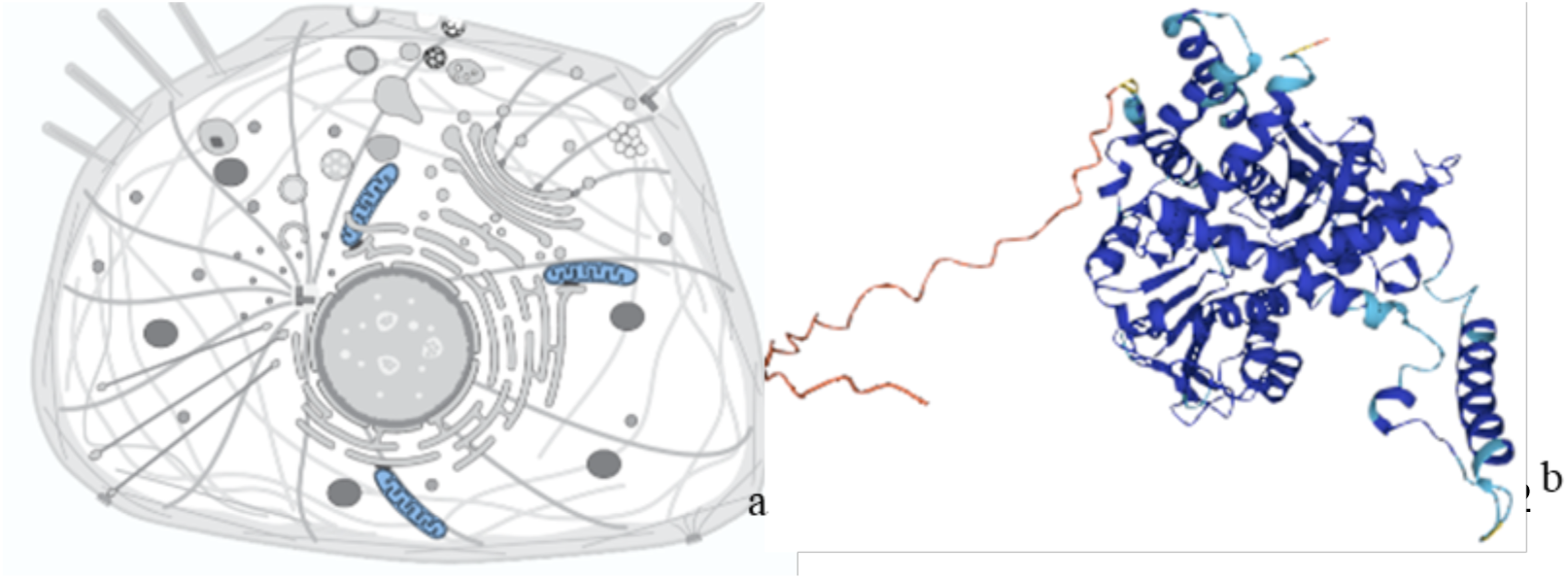
a) Localization of the glutamate dehydrogenase, mitochondrial isoform X2, and b) Tridimensional structure. Source: UniProt, 2023.

From a nutraceutical and functional perspective, the identified proteins offer potential due to their antioxidant activities, metabolic control, and cellular regulation, since they may have food applications with health benefits according to various studies that support these effects. For example, the enzyme aldose reductase has been documented as important in sugar metabolism and plays therapeutic roles in diabetes control (Hernández et al., 2011). Likewise, glutamate dehydrogenase (GDH) is an important enzyme in amino acid metabolism that provides energy and gluconeogenic substrates (Gaspar et al., 2018; Smith et al., 2019), whereas heat shock protein 60 A (HSP60) has therapeutic potential in cardiovascular diseases (Krishnan et al., 2021).

The proteins identified during the developmental stages of queen bees reveal a complex network of molecular mechanisms regulating several processes: from hormonal regulation controlling vitellogenin synthesis to the synthesis of proteins involved in cellular protection such as HSP60 A and thioredoxin reductase; each of these proteins plays a crucial role in metamorphosis. Ubiquitination enzymes and proteins involved in cellular metabolism, such as the electron transfer flavoprotein and aldose reductase, ensure that cells can respond to stress and maintain the energy necessary to complete development, highlighting the biological sophistication of queen bees during their transition to the adult stage.

## Conclusions

The proteomic analysis of *Apis mellifera* L. pupae reveals a complex protein profile that varies according to their developmental stage and is expressed in fundamental aspects related to metabolism, the response to oxidative stress, and cellular regulation. The proteins identified in this study suggest significant nutraceutical and functional potential due to their diverse biological activities and beneficial properties for health.

This provides new perspectives on the potential consumption of queen bee pupae as a food and opens possibilities for the development of nutritional supplements. Such applications could contribute significantly to food security and human health by offering nutritious and functional alternatives based on this apicultural product.

## Data availability statement

The data (Supplement Proteome paper and Abstract Image) that support the findings of this study are openly available in figshare at https://doi.org/10.6084/m9.figshare.31549972

## Graphical Abstract or Cover Image Note

The graphical abstract (cover image) included in this submission was not generated by artificial intelligence.

## Authors contribution (Credit taxonomy)

Conceptualization: R. De la Rosa-Santamaría Metodology: R. De la Rosa-Santamaría, I. López-Rosas

Formal Analysis: R. De la Rosa-Santamaría, I. López-Rosas, L. M. Vargas-Villamil,. Investigation: R. De la Rosa-Santamaría, I. López-Rosas, D. G. Ruiz-Pérez

Data curation: R. De la Rosa-Santamaría, I. López-Rosas, D.G. Ruiz-Pérez Writing–original draft: R. De la Rosa-Santamaría, I. López-Rosas

Writing – revision and edition: I. López-Rosas, S. Cadena-Villegas, Jenner Rodas-Trejo,

L.M. Vargas-Villamil. Francisco Izquierdo-Reyes Supervision: R. De la Rosa-Santamaría

